# Ionotropic glutamate receptors in the retina – a bioinformatical meta-analysis

**DOI:** 10.1101/2021.11.27.470202

**Authors:** Bianca Pircher, Thomas Pircher, Andreas Feigenspan

## Abstract

Glutamate is an essential neurotransmitter for signal processing in the vertical pathway of the mammalian retina, where it is involved in the distribution of visual information into several parallel channels. The excitatory effects of glutamate are mediated by AMPA-, kainate-, and NMDA-type ionotropic glutamate receptors (iGluRs). The expression patterns of these receptors in the vertebrate retina have been investigated so far with mainly immunocytochemical, in-situ hybridization, and electrophysiological/pharmacological techniques. Here, we have used scRNA sequencing data from chicken, mouse, macaque, and human retina to describe and compare the profile of iGluR expression in major retinal cell types across species. Our results suggest that major retinal cell types each express a unique set of iGluRs with substantial differences between non-mammalian and mammalian retinae. Expression of iGluRs has been investigated in more detail for amacrine and bipolar cell types of the human retina, each showing minor variations of a common pattern. The differential expression of iGluRs is likely to convey unique signal processing properties to individual elements of the retinal circuitry.

## Introduction

L-glutamate is a ubiquitous neurotransmitter in the vertebrate central nervous system, and it also plays a decisive role in the synaptic circuitry of the retina [1]. The broad diversity of postsynaptic glutamate receptors is key for transmission of intricate signal properties, especially in sensory systems like the retina.

In general, glutamate mediates its actions by activating members of the ionotropic (iGluR) and metabotropic glutamate receptor (mGluR) families; however, only iGluRs will be addressed in this study. Based on their pharmacological and electrophysiological properties, iGluRs can be classified into N-methyl-D-aspartate (NMDA) and non-NMDA receptors, which are further divided into AMPA (*α*-amino-3-hydroxy-5-methyl-4-isoxazolepropionic acid; GluA1-GluA4) and kainate (2-carboxy-3-carboxymethyl-4-isopropenyle-pyrrolidine; GluK1-GluK5) receptors [2]. All iGluRs are homo- or heteromeric protein complexes composed of four large subunits that form a non-selective cation channel. In addition to the pore-forming transmembrane domain (TMD), glutamate receptor subunits consist of three additional distinct and semiautonomous domains: the extracellular amino-terminal domain (ATD), also referred as the N-terminal domain (NTD), the extracellular ligand-binding domain (LBD), and the intracellular carboxyl-terminal domain (CTD) [2], [3]. Figure 1 shows the schematic domain structure of an AMPA receptor subunit.

**Figure 1.**
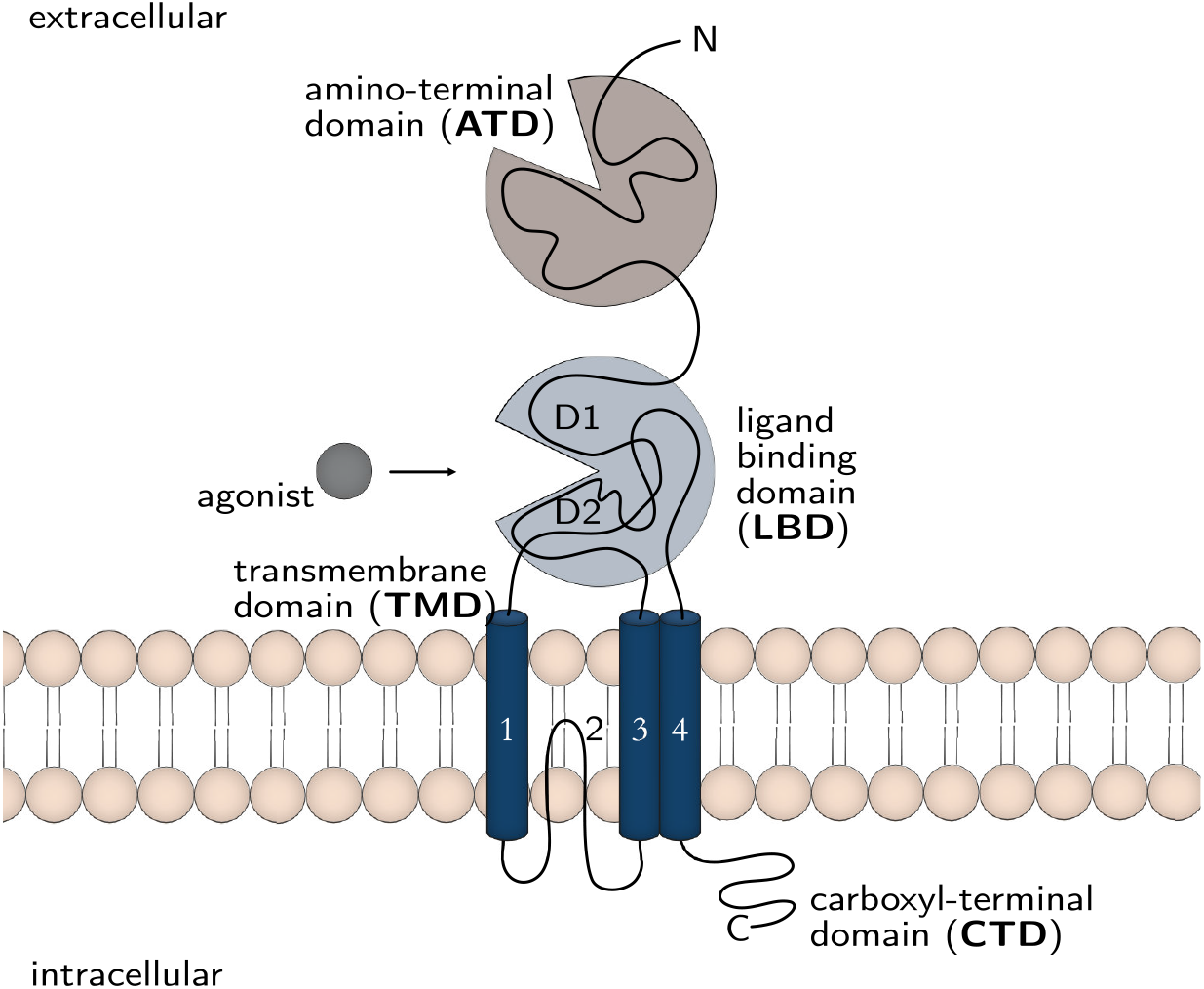
Schematic illustration of an ionotropic glutamate receptor subunit. The template for this illustration was the AMPA receptor GluA2. Glutamate receptors have a modular structure composed of the amino-terminal (ATD), the ligand binding (LBD), the transmembrane (TMD) and the carboxyl-terminal domain (CTD). The TMD, which forms the ion channel pore, is built up of three membrane-spanning helices (1, 3 and 4) and a re-entrant loop (2).

The extracellular components encompass the largest part of the receptor subunit, and they are arranged as a dimer of dimers. Therefore, they display a 2-fold symmetry, whereas the TMD shows a 4-fold symmetry [4]. The TMD is composed of three helical elements (M1,M3, and M4), and a membrane re-entrant loop (M2) that forms the ion-selective filter of the pore [3]. The overall structure of the iGluR channel pore is similar to an inverted potassium channel pore with M2 entering the membrane from the extracellular side [5], [6], [7].

NMDA-receptors (NMDARs) can be distinguished from other iGluRs, because the channels are blocked by extracellular magnesium (Mg^2+^) at physiological resting membrane potentials, and they open only upon agonist binding in combination with simultaneous depolarization, which expels Mg^2+^ from the channel pore. For an efficient activation, the co-agonist glycine is required in addition to glutamate [8]. The heteromeric complexes are assembled from a repertoire of three major subunits: GluN1, GluN2, and the one discovered latest, GluN3, which are encoded by seven different genes [9], [10] (see Table 1a). In addition, numerous splicing variants are known, especially for GluN1. A functional NMDAR requires co-expression of at least one GluN1 and one GluN2 subunits [10]; thus, two GluN1 and two GluN2 subtypes frequently assemble to form a functional NMDAR [11].

**Table 1.**
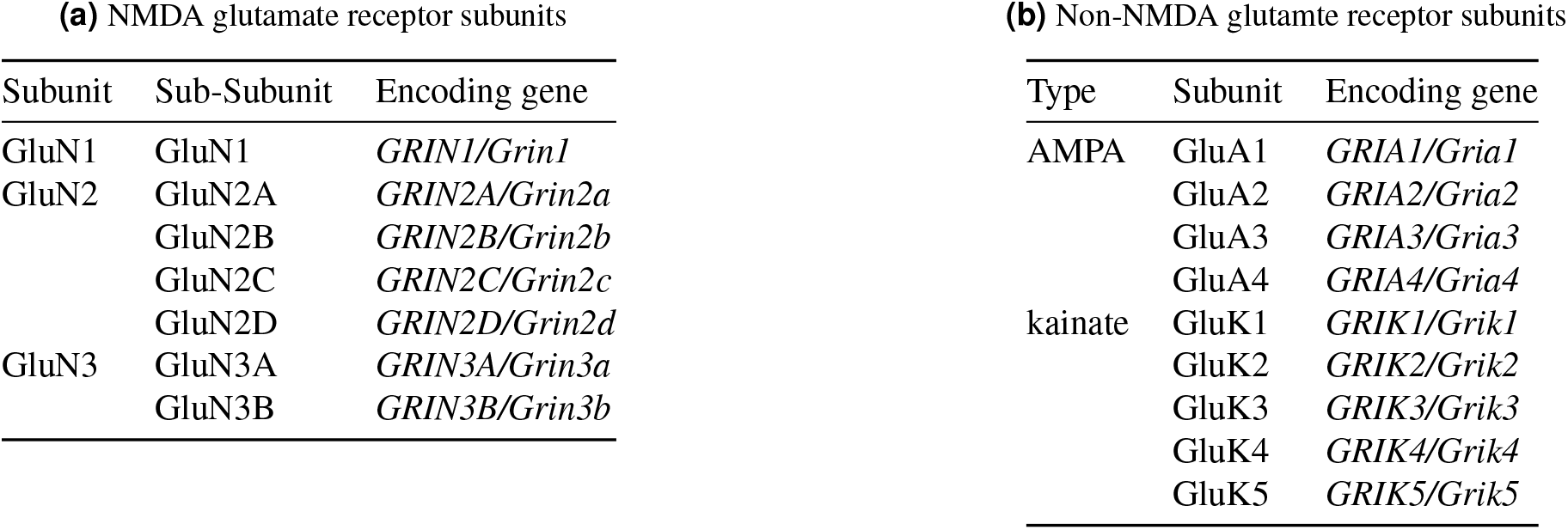
Ionotropic glutamate receptors with their corresponding subunits and encoding genes.

In the central nervous system, NMDARs are of particular interest, since they are involved in synaptic plasticity, neurodegeneration, depression, and chronic pain [8], [2]. Although NMDARs are present in the vertebrate retina, their expression profile has remained a matter of controverse. While immunocytochemical or *in situ* hybridization studies argued for the expression of NMDARs subunits in various retinal cell types [12], [13], electrophysiological approaches were often not able to support these findings [14]. This is particularly evident for horizontal cells and bipolar cells. In horizontal cells, electrophysiological studies rejected the presence of NMDARs [15], [16], whereas immunocytochemical evidence for expression of GluN1 and GluN2 was obtained [17], [18]. These differences, however, could be due to species-specific (carp, rabbit, rat) expression of NMDARs. Very similar findings were reported for bipolar cells [14]. With respect to expression of NMDARs in the inner nuclear layer and ganglion cell layer, immunocytochemistry and electrophysiology are in better agreement.

In several of the more than sixty different types of retinal amacrine cells [19], immunolabeling for NMDAR subunits [20], [21] as well as distinct electrophysiological responses [22], [23], [24] were observed [25]. Retinal ganglion cells have been shown to contain the subunits GluN1 and GluN2, which apperently assemble into functional NMDARs [26].

AMPA receptors, which mediate fast excitatory synaptic transmission, are composed of four GluA1-4 subunits, either as homo- or hetero-oligmers [27], [2]. The subunits are encoded by separate genes termed *Gria1-4*. Several studies indicate that diverse heteromeric AMPA receptor subtypes are arranged in a region- and cell-type specific manner [28], [29], [2]. In addition, the expression of all subtypes is activity-dependent, which emphasizes their role for synaptic plasticity [2]. The variable assembly of four different subunits in combination with RNA editing and alternative splicing leads to combinatorial diversity and distinct functional AMPA receptor properties, which is reflected in diverse channel kinetics and conductances [28].

Kainate (KA) receptors are expressed both pre- and postsynaptically throughout the entire central nervous system, where they modulate transmitter release and contribute to synaptic transmission, respectively [30], [31], [32], [33]. The encoding genes are termed *Grik1-5*. In contrast to AMPA receptors, which are rather well-characterized with respect to their physiological properties and their role for synaptic transmission, less is known about KA receptors [34], [33], largely due to the lack of selective pharmacological tools [34]. KA receptor subunits can form homo- and heteromers composed of the subunits GluK1 to GluK5 (see also Table 1b), some of which are susceptible to alternative splicing and mRNA editing [31]. It is suspected that receptors containing GluK4 and GluK5 require coexpression of GluK1 to GluK3 for proper functionality [2]; however, general rules governing the subunit assembly of KA receptors have not been described yet [35]. GluK1-3 subunits apparently show low-affinity for glutamate binding, whereas GluK4 and GluK5 are high-affinity subunits [33]. So far it is not known whether the subcellular localization of KA receptors is relatated to their subunit composition, or vice versa [33].

The localization of AMPA and KA receptors in the vertebrate retina has been extensively studied immunocytochemically in several species [36], [37], [38], [39], [40], [41], [42], [43], [44], [45], [46], and these results are supported by electrophysiological investigations. Bipolar cells have been shown to express both AMPA and kainate receptors [47]. The majority of amacrine cells contain both receptor types, whereas some cell types exclusively express either AMPA or KA receptors [23]. Of particular interest is GluA2, as it is involved in the regulation of dopaminergic amacrine cells by light [48]. *In situ* hybridization and immunostaining carried out in retinal ganglion cells provide evidence for expression of all subtypes of both AMPA and KA receptors [27]. The situation in horizontal cells is more complex and remains controversial [49], [50]. While some electrophysiological studies have found evidence for both receptor types ([51], [52], [49]), a recent study has excluded the involvement of kainate receptors in signal processing of horizontal cells [53]. Immunocytochemical data indicate the presence of both AMPA and KA receptors [20], [54], since GluK2 and GluK3 (formerly GluR6 and GluR7) have been detected in several mammalian species (rat: [13]; cat: [55]; [56]; primate: [57]). AMPA-receptors, namely GluA2, GluA4, and possibly GluA3 were detected immunocytochemically in the mouse retina [54].

Our current knowledge concerning the expression of iGluRs in the mammalian retina relies largely on immunocytochemical and pharmacological evidence, which is limited by the specificity of antibodies and the availability/affinity of subtype-specific drugs, respectively. However, scRNA-sequencing is a promising complementary approach, which allows the analysis of RNA expression of nearly countless genes in different cell types and even across species at the same time. Here, we provide a comprehensive overview on the subtype-specific expression of NMDARs and non-NMDARs in the vertebrate retina using data sets from four different species (mouse, human, macaque and chicken).

## Results

To investigate the expression of iGuR subtypes in photoreceptors, horizontal cells, bipolar cells, amacrine cells and retinal ganglion cells, we analyzed four freely available scRNA-seq data sets of different vertebrate species.

First, as one of the most important model organisms in eye research, we analyzed a scRNA-seq data set of the mouse retina [58]. The results are depicted in Figure 2.

**Figure 2.**
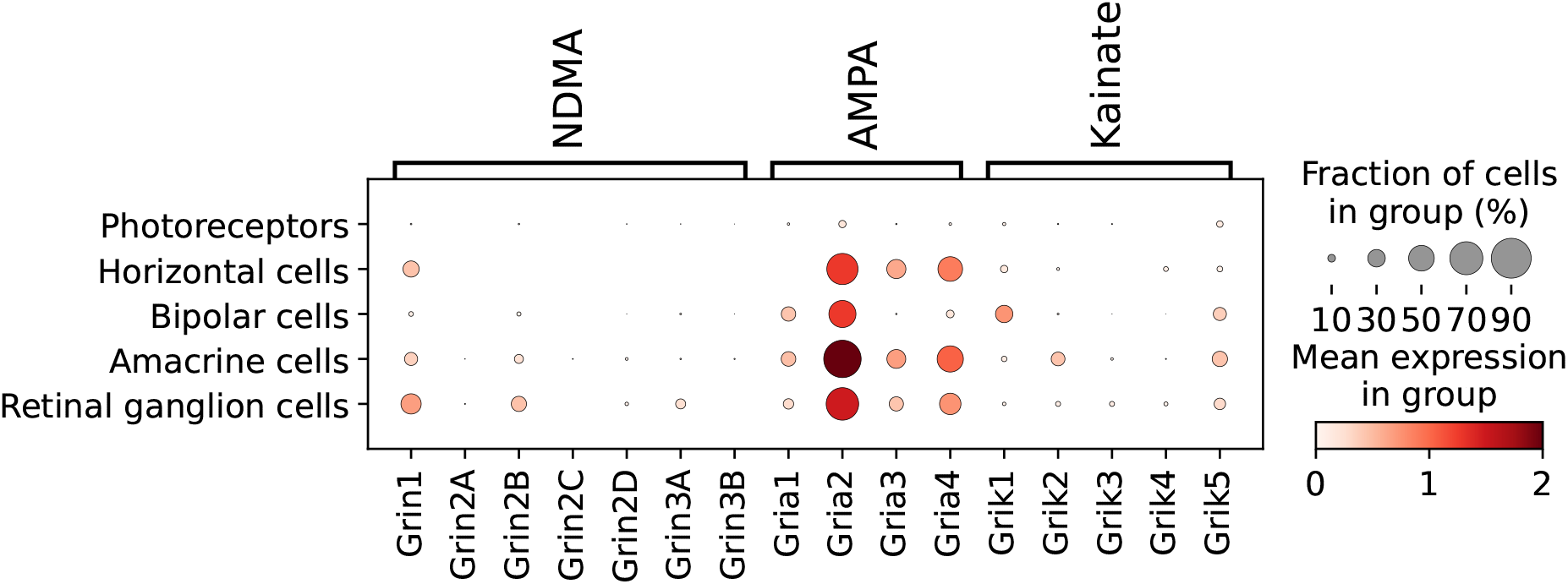
Expression of different glutamate receptor-encoding genes in major cell types of the mouse retina. Visualized data set is based on scRNA-seq data set of Macosko et al. [58].

At first glance, the pronounced expression of the AMPA subtypes is obvious. Especially *Gria2*, the gene encoding for the GluA2 subunit, is highly expressed in all cell types, except photoreceptors, which apparently do not contain iGluRs. *Gria3* and *Gria4* are expressed in horizontal, amacrine, and retinal ganglion cells. In contrast, the composition of iGluRs in bipolar cells is different, since they do not express NMDA receptors, but both AMPA and KA receptors. Based on this data set, bipolar cells presumably contain GluA1, GluA2, GluK1, and possibly GluK5. Horizontal cells express AMPA receptor subunits GluA2, GluA3 and GluA4, but no kainate receptors. In contrast to previous results, which rule out expression of NMDA receptors in horizontal cells [15], [16], this data set suggests the existence of the NMDA receptor subunit GluN1. The receptor composition of amacrine and retinal ganglion cells seems to be quite similar. Both are apparently equipped with GluA1, GluA2, GluA3, GluA4, GluK5, and GluN1. Retinal ganglion cells additionally express *Grin2B*, econding for GluN2, and *Grin3A*, encoding for GluN3, whereas amacrine cells show the additional expression of GluK2.

Next, we analyzed the scRNA-seq data set from the human retina [59]. The expression of AMPA receptor subunits across cell types appears rather similar to the pattern observed in the mouse retina (Fig. 3), although the expression of *GRIA2* is not as pronounced. Concerning kainate receptors, only GluK1 is expressed in bipolar cells, and GluK5 might be present in ganglion cells. Expression of kainate receptors could not be verified in other cell types of the human retina. NMDA receptors are predominantly present in retinal ganglion cells with an emphasis on GluN1, GluN2A and GluN2B; however, GluN1 is absent in horizontal cells. Expression of *GRIN1* in amacrine cells appears likely.

**Figure 3.**
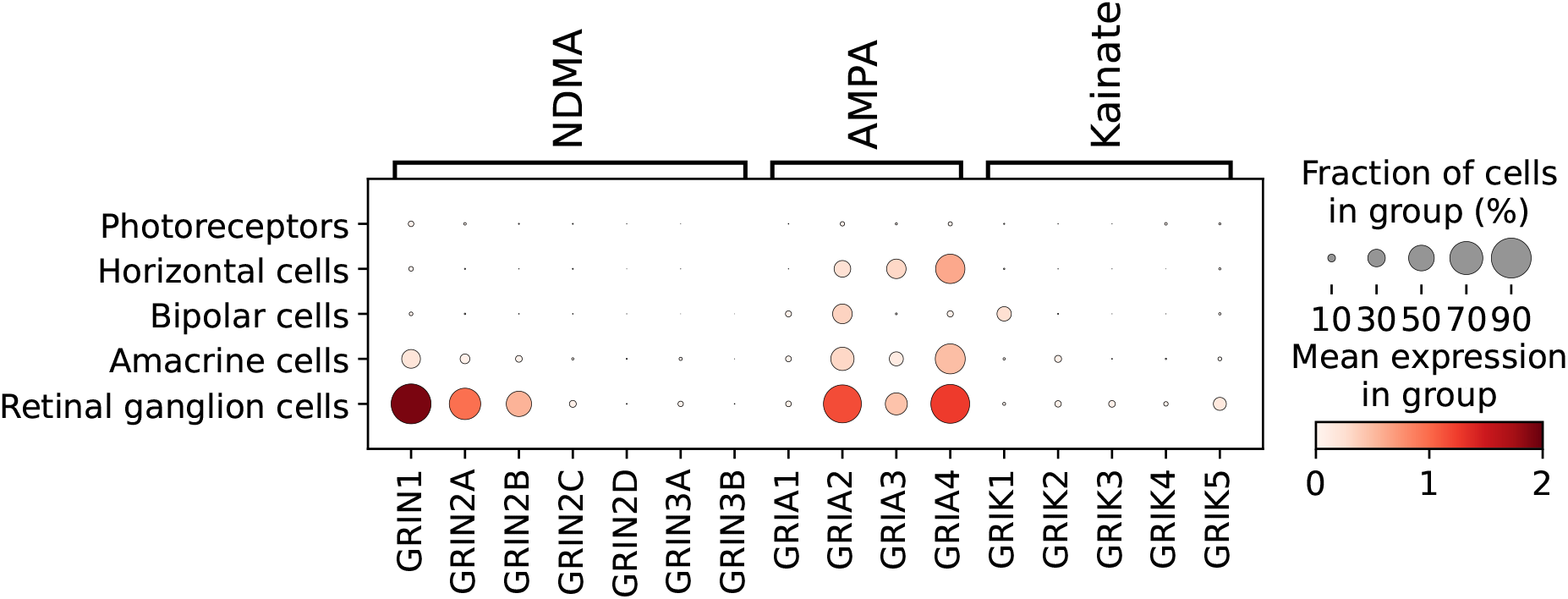
Expression of different glutamate receptor encoding genes in major cell types of the human retina. Visualized data set is based on scRNA-seq data set of Yan et al. [59].

Compared to the human retina, the macaque retina shows a nearly identical pattern of AMPA receptor distribution, and also the prevalence of NMDA receptor encoding genes in retinal ganglion cells is quite similar (Fig. 4). The only notable difference is the absence of *Grin2B* expression in ganglion cells. *Grin2D*, *Grik1* and *Grik5* are entirely missing in this data set. We assume that these subunits were so scarce that they were not deposited in the data set as existing genes. Parallels between bipolar cells that express *Grik1* and retinal ganglion cells that exhibit *Grik5* in both data sets thus cannot be drawn.

**Figure 4.**
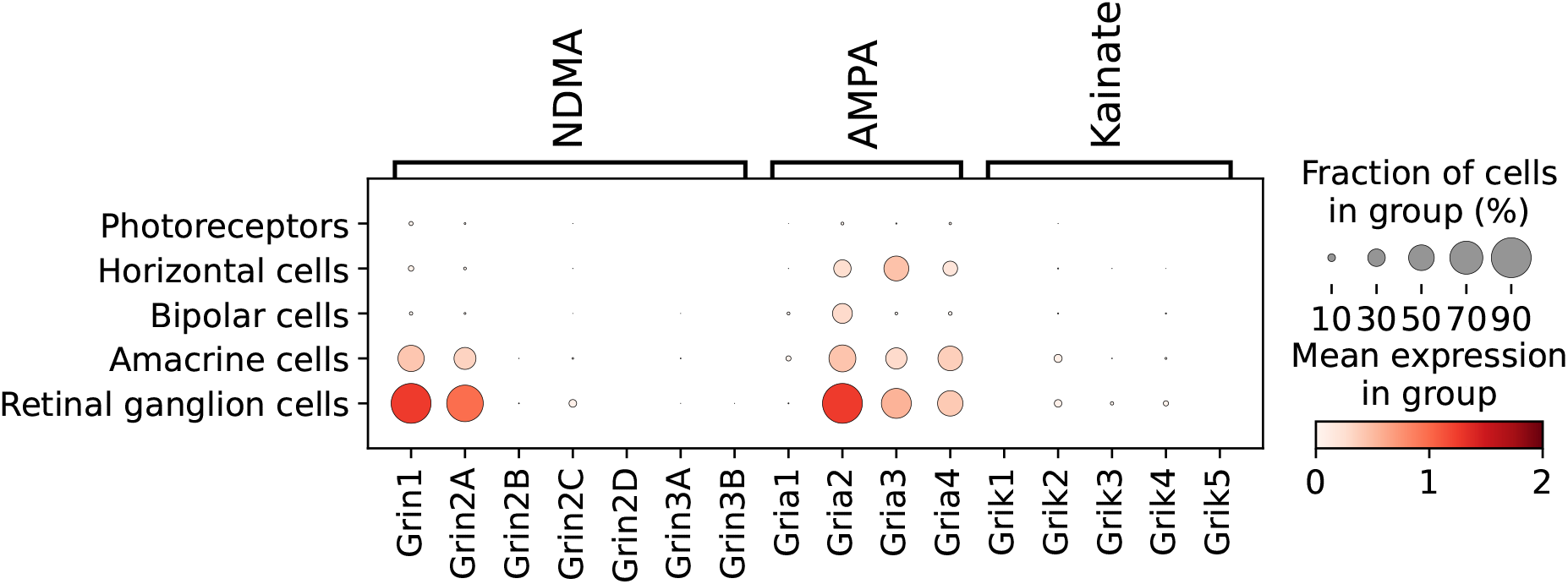
Expression of different glutamate receptor encoding genes in major cell types of the primate retina. Visualized data set is based on scRNA-seq data set of Peng et al. [60].

In contrast to the mammalian data sets, distinct differences in gene expression can be observed for the chicken retina (Fig. 5). As the only non-mammalian species in this analysis, differences in the configuration of NMDA receptors are particularly apparent. Amacrine cells and ganglion cells, as described in the other species, but also horizontal cells and bipolar cells express different subunits of NMDA receptors. The most abundant of these is the GluN1 subtype. Bipolar cells, however, seem to express only GluN1 and the kainate subtype GluK1, whereas AMPA receptor encoding gene could not be detected. Also GluA2, predominant in the mammalian retina, is differently expressed in the chicken. In horizontal cells and bipolar cells there is almost no expression of *Gria2*, whereas it is present in photoreceptors, which also show a low-level expression of *Grik1* and *Grik3*. Similar to the mammalian data sets, the KA receptor subtypes *Grik1-3* are also expressed by amacrine cells and retinal ganglion cells.

**Figure 5.**
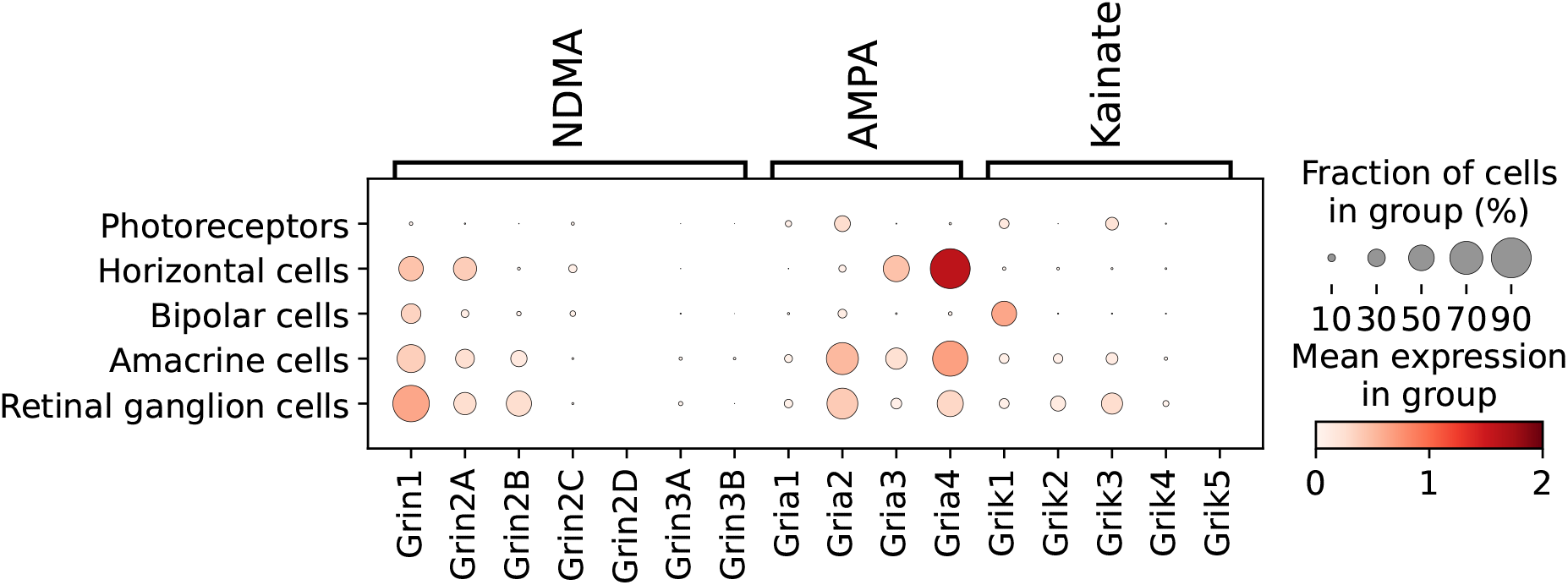
Expression of glutamate receptor encoding genes in major cell types of the chicken retina. Visualized data set is based on scRNA-seq data set of Yamagata et al. [61].

In the previous graphs, we showed differences in the expression patterns of iGluRs in the major retinal cell types of various vertebrate species. However, it should be noted that some of these major cell types comprise numerous, partly very specific subtypes, which differ in their connectivity, morphology, and neurotransmitter content. For example, amacrine cells have been proposed to contain more than 60 different subtypes by a comprehensive bioinformatical approach [19]. In order to emphasize this diversity and to gain a more complete picture of the different amacrine cell subtypes, we extended our analysis to this particular data set [19]. Amacrine cells of the mouse retina were grouped according to their neurotransmitter content into GABAergic, glycinergic, cholinergic, dopaminergic and glutamatergic cells (see Methods for further details). As shown in Figure 6, these amacrine cell subtypes display a rather uniform distribution of AMPA and kainate receptors, which is mostly in accordance to the findings of Macosko et al. [58] (Fig. 2). The expression of *Gria3* appears slightly more prominent in glutamatergic amacrine cells. The major difference between the two data sets of the mouse retina relates to the expression of *Grin2a*. In the data set of Macosko et al. [58] this gene is not expressed at all in amacrine cells, whereas in the data set of Yan et al. [19], *Grin2a* is clearly present in glutamatergic amacrine cells. *Grin2a* appears not to be expressed in cholinergic amacrine cells, which show highest expression of *Grin2b* compared to other amacrine cell subtypes. With the exception of *Grik5*, which appears at similar levels in all amacrine cell subtypes, the expression of other kainate receptors seems marginal.

**Figure 6.**
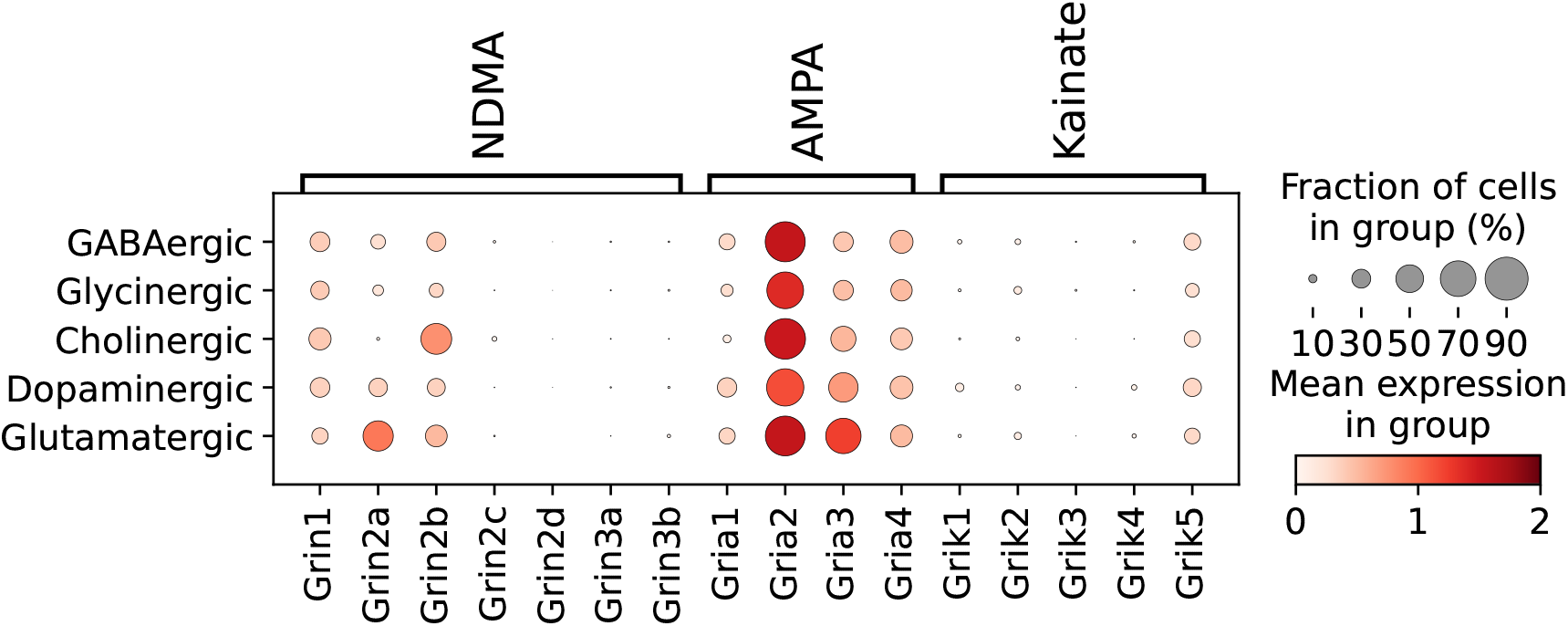
Expression of glutamate receptor encoding genes in different types of amacrine cells grouped by their major neurotransmitter. Visualized data set is based on mouse scRNA-seq data set of Yan et al. [19]

Various types of bipolar cell contribute decisively to vertical information processing in the retina. For the current investigation we used the human data set of Yan et al. [59], since the authors have already performed a comprehensive clustering into 12 known and well characterized bipolar cell subtypes. To compare the expression of iGluRs in the major bipolar cell types, we grouped the subtypes into ON cone bipolar cells, OFF cone bipolar cells, as well as rod bipolar cells. The expression pattern of glutamate receptor encoding genes in these different groups of bipolar cells is visualized in Figure 7. Interestingly, only OFF cone bipolar cells contain *GRIK1*, which encodes the kainate receptor GluK1. Neither ON cone bipolar nor rod bipolar cells show expression of *GRIK1* or of any other kainate receptor. *GRIA2* appears evenly distributed between the three major subtypes with the exception of DB3b, which only shows low-level expression of *GRIA2*. In fact, DB4 does not show any expression of ionotropic glutamate receptor subunits.

**Figure 7.**
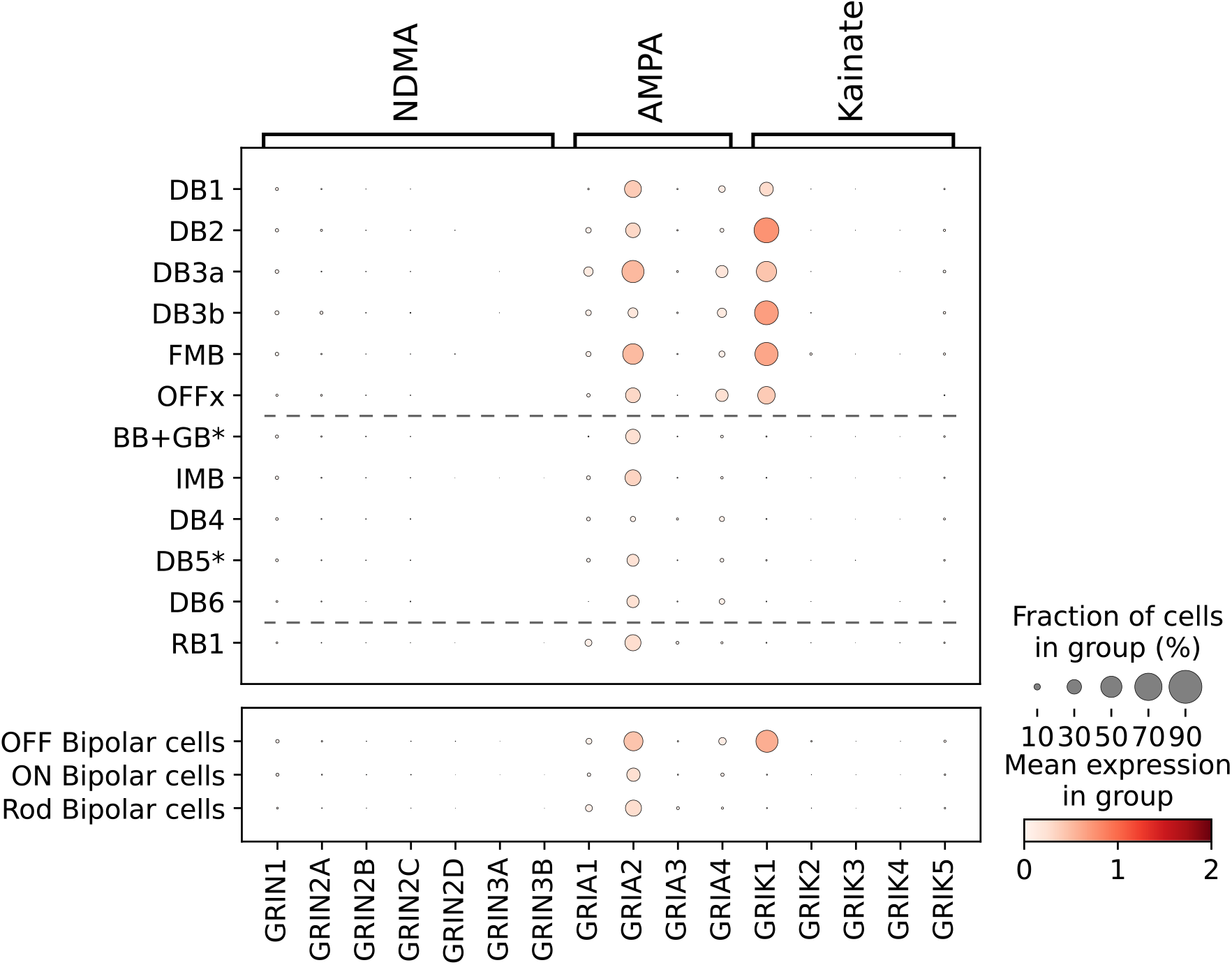
Expression of glutamate receptor encoding genes in different types of bipolar cell (top) and grouped into OFF, ON and Rod bipolar cells (bottom). Dotted lines in the top panel delineate the border between OFF and ON cone bipolar cells and between ON cone bipolar cells and rod bipolar cells, respectively. Visualized data set is based on human scRNA-seq data set of Yan et al. [59].

## Discussion

In the present study, we investigated the expression of ionotropic glutamate receptors in the retina of several mammalian and one avian species using for the first time a bioinformatical approach. In contrast to previously published immmunocytochemical and pharmacological studies, which are often impeded by the specificity of antibodies and antagonists available, the analysis of cell type-specific gene expression promises an unbiased and in-depth account of glutamate receptor distribution in the vertebrate retina.

Perhaps most striking is the difference between the mammalian species and the only non-mammalian representative, the chicken. Whereas in the chicken most retinal cell types are well equipped with NMDA receptors, this receptor type is confined to amacrine cells and ganglion cells in the mammalian retina. In the chicken retina, all cells except photoreceptors express at least one NMDA receptor subunit, predominantly *Grin1*. In addition, each retinal cell type appears to express a distinct combination of iGluRs. Horizontal cells, for example, do not contain *Gria2*, but *Gria3* and *Gria4*, as well as two genes encoding for NMDA receptor subunits (*Grin1* and *Grin2A*). Bipolar cells apparently are not equipped with AMPA receptor subunits. Both observations are in agreement with an immunocytochemical study in the developing chick retina [62], where the authors report the absence of labeling of any AMPA receptor subunit in bipolar cells and also a distinct staining of GluA4 in horizontal cells.

In contrast, immunocytochemical evidence in the pigeon retina indicates expression of several AMPA receptor subtypes in bipolar cells [46]. OFF bipolar cells were reported to express GluA1 through GluA4, GluK1, GluK2, and GluK4 as well as GluN1 and GluN2A. These results suggest major differences in the expression of iGluRs in mammalian compared to avian retina in addition to differences within the avian taxon itself. These discrepancies might relate to differential signal processing in these vertebrate classes.

Similar considerations might also apply to the subclass of placental mammals even if the pattern of AMPA receptor subunit-encoding genes seems to be conserved to some extent in the species under investigation. Horizontal cells, amacrine cells, and retinal ganglion cells express *Gria2*, *Gria3* and *Gria4*, whereas bipolar cells express *Gria2* only. This pattern was observed in the mouse, human, and in the macaque retina. In contrast, expression of *Gria1* appears more heterogeneous, with detectable contents only in the mouse retina. A direct comparison of major cell types, however, is limited, since species differ considerably in cell-specific subtypes, which are likely to precisely tune information processing within the retinal circuitry.

### Photoreceptors

Our bioinformatic approach indicates marginal, if any, expression of iGluRs in photoreceptors of all mammalian species tested, which is in agreement with the results of Brandstätter et al. in the rat retina [13]. In contrast, Haumann et al. [33] report a close association of GluK5 with the ribbon of mouse rod but not cone photoreceptors as well as an alteration of presynaptic structure in GluK5-deficient mice. However, the existence of other GluK subtypes necessary for a fully functional kainate receptor [2] have not be shown. Although we observed an exceedingly small spot representing *Grik5* in the mouse retina, we could not detect expression of other kainate receptor encoding genes in all mammalian species investigated. Therefore, a possible autoregulatory or structural function of GluK5 in rod spherules requires further evidence.

### Horizontal cells

The expression pattern of iGluRs in mammalian horizontal cells has been a continuing matter of debate. Although most immunocytochemical and electrophysiological studies suggest the existence of kainate receptors in horizontal cells [20], [54], [51], [52], [49], this conclusion has most recently been rejected [53]. The bioinformatical approach presented here also argues against the expression of kainate receptors in horizontal cells in mouse, primate, and human retina. The expression of AMPA receptors, however, appears to be in agreement with previous experimental results [54]. The genes *Gria2-4*, encoding for GluA2-GluA4, show distinct expression in mammalian horizontal cells. The presence of NMDA receptors, on the other hand, again is controversial. Whereas electrophysiological experiments showed no evidence for the presence of NMDA receptors in horizontal cells [15], [16], imunocytochemical studies suggested the expression of *Grin1* and *Grin2a* [17], [18]). We report a distinct expression of these two receptor subunits in the mouse (*Grin1*) and in the chicken (*Grin1* and *Grin2*). However, a functional role of NMDA receptors in horizontal cells remains to be determined.

### Bipolar cells

The expression of AMPA and kainate receptors in OFF bipolar cells has been shown previously with an immunocytochemical approach [41], [57], [63], and our bioinformatical results also indicate expression of GluA2, GluK1, and possibly GluA4. However, recent studies including patch-clamp measurements from bipolar cells of macaque and mouse retina rejected the notion of functional AMPA receptors in OFF bipolar cells [64], [65]. Puthussery et al. [64] showed that OFF bipolar cells of the primate retina process their synaptic input exclusively via kainate receptors. The authors also claimed that GluK1, which in our study appears to be expressed in all types of OFF bipolar cell, is confined to DB2 and DB3b cells, whereas other OFF cone bipolar cells lack GluK1 and presumably express GluK2 or GluK3. Even though it is evident from our data that DB1 and Db3a express GluK1, the expression of GluK2 and GluK3 is completely absent from these cells. Moreover, this observation can be extended to other species investigated. Only the mouse showed some expression of *Grik5* in bipolar cells. Other studies in the mouse and in the ground squirrel retina stated that each morphological type of OFF cone bipolar cell receives its signal through a unique combination of AMPA and kainate receptors as a prerequisite for diversity in the temporal domain [47], [45], [66], [67]. Additionally, Puller et al. [45] reported that AMPA and kainate receptors are not localized at the same synaptic sites in type 3b and type 4 Off cone bipolar cells.

The bioinformatic approach clearly suggests that only OFF but not ON cone bipolar cells of the human retina express kainate receptors, i.e. GluK1. In general, metabotropic glutamate receptor type 6 (mGluR6) is supposed to mediate the sign-converting signal in ON cone bipolar cells [68], [69]. However, immunocytochemical evidence for the presence of AMPA receptors (especially GluA2) in rod and cone ON bipolar cells has been reported [63],[70], which is consistent with our results.

In summary, the precise pattern of ionotropic glutamate receptor expression in the different bipolar cell types remains elusive.

### Amacrine cells

Previous immunocytochemical studies suggest the expression of several subtypes of AMPA, kainate, and NMDA receptors in amacrine cells of the mammalian retina [37], [36], [12], [21], [71], [38], which is generally confirmed by the data presented here. Interestingly, amacrine cell types as distinguished by their neurotransmitter content, display a rather uniform expression of iGluRs. *Gria2*, encoding for the GluA2 receptor subtype, was found in all amacrine cells, and the activation of this Ca^2+^-permeable AMPA receptor is necessary for amacrine cell plasticity [72]. The ubiquitous presence of GluA2 in all amacrine cell types appears to be inconsistent with the findings of Dumitrescu et al. [23], who reported a differential expression of AMPA and kainate receptors in amacrine cells of the mouse retina. In a human data set, we detected the rather distinct expression of GluK5, which alone is not sufficient to assemble into functional kainate receptor channels. However, we cannot rule out low-level expression of other kainate receptor subtypes, especially GluK2, in the mouse retina. Therefore, species-specific differences might be responsible for these conflicting results.

The AII amacrine cell is of key importance for funnelling scotopic signals into ON and OFF channels in the inner plexiform layer. AII amacrine cells are glycinergic neurons that have been shown electrophysiologically to contain both NMDA and non-NMDA receptors [73] (but see [74]). Pharmacological experiments and paired recordings have provided evidence for AMPA receptors, but not kainate receptors in amacrine cells of the rat retina [75], [76]. Our data show a preponderance of AMPA receptor-encoding genes in glycinergic and also in other amacrine cell subtypes grouped by neurotransmitter content. According to this data set, the equipment with iGluRs differs only slightly between these major subgroups, with apparently higher expression of GluN2B in cholinergic and GluN2A in glutamatergic amacrine cells.

The bioinformatical results indicate pronounced expression of AMPA and NMDA receptors, but allow little, if any conclusions on the involvement of kainate receptors. A detailed investigation of cell-type specific expression of iGluRs in amacrine cells is necessary for a profound understanding of signal processing in the inner retina. Given the plethora of around 60 different amacrine cell types, our classification according to neurotransmitter content serves only as a very superficial guidance.

### The open question of functional receptors

Even if the presence of an iGluR subtype is confirmed in a specific retinal cell type, one always has to consider the requirements for the assembly of functional glutamate receptor channels. As already mentioned, functional NMDA receptors require co-expression of at least one GluN1 and one GluN2 subtype [10]. Therefore, the exclusive expression of *Grin1* in mouse horizontal cells does not suffice for a fully functional NMDA receptor. This might explain the failure of electrophysiological studies to detect NMDA receptors in this cell type [15], [16], whereas some immunocytochemical evidence was found [17], [18]. The same reasoning applies to the expression of *Grin1* in bipolar cells of the chicken retina.

The situation for GluA2-containing AMPA receptor is different. Here, the homomeric assembly of GluA2 into functional receptors has been shown currently, since this subunit is subject to RNA editing in the selectivity filter [77]. Therefore, the seemingly exclusive expression of *Gria2* in bipolar cells of the mammalian species does not exclude the presence of a functional glutamate receptor.

We also observed different kainate receptor subtypes in the expression dot plots. In particular GluK1, often exclusively expressed in bipolar cells, suggests the formation of a homomeric kainate receptor. As it was stated, GluK1-3 can indeed form functional homomeric receptors [78]. Only GluK4 and GluK5 subunits are not able to assemble into functional homomeric receptors, as they require coexpression with GluK1 through GluK3 subunits. GluK2/K5 receptors, being the most abundant combination in the CNS [79], seem to be mostly absent in the retina. Only in the data sets of Macosko et al. [58] and Yan et al. [59] both subunits might be expressed in amacrine cells.

Finally, it should be noted that no valid statement can be made about the functionality of a glutamate receptor channel based solely on scRNA seq expression data. Therefore, physiological and immunocytochemical evidence is always necessary to substantiate the functional role of iGluRs within the retinal circuitry. However, the analysis of scRNA data establishes the possibility of a comprehensive account of a cells’ channel inventory, allowing direct comparison between cell types of a given species and also between species of different evolutionary distance. Therefore, scRNA data sets should be expanded, especially for the important model organisms zebrafish and salamander, which are both extensively used in retinal research.

## Methods

To investigate the expression of ionotropic glutamate receptor subunits in the major classes of retinal cell types, we used data sets from four different species (mouse: (GSE63473) [58], human: (GSE148077) [59], macaque: (GSE118480) [60], and chicken: (GSE159107) [61]).

The data set of Macosko et al. [58] provides single cell sequencing results for the complete mouse retina and its distinct cell types. For the analysis, we used the clustering as published and available at https://singlecell.broadinstitute.org/single_cell/study/SCP7/drop-seq. We extracted cells assigned as retinal ganglion cells, amacrine cells, horizontal cells, bipolar cells, and photoreceptors. In the latter case, rods and cones were combined. All retinal cells were tested for their mean expression of genes encoding for NMDA (*Grin1, Grin2a, Grin2b, Grin2c, Grin2d, Grin3a, Grin3b*), AMPA (*Gria1, Gria2, Gria3, Gria4*) and kainate receptor subunits (*Grik1, Grik2, Grik3, Grik4, Grik5*). Other cell types like Müller glial cells were not extracted nor included in the analysis.

The same procedure was conducted with the human retinal data set of Yan et al. [59] (https://singlecell.broadinstitute.org/single_cell/study/SCP839) and with the chicken data from Yamagata et al. [61] (https://singlecell.broadinstitute.org/single_cell/study/SCP1159/). As the clustered data was deposited differently on the single cell portal (https://singlecell.broadinstitute.org/single_cell/study/SCP212) by the authors, the procedure to analyse the macaque data set was slightly different. In their work the authors compared the gene expression in retinal cell types of the fovea and of cells in the periphery. Therefore, we downloaded both expression files for amacrine, bipolar, horizontal and retinal ganglion cells and combined them as a repository for the entire retina. Unfortunately, at the time of downloading the data (2020-08-20), the data set of peripheral photoreceptors was damaged and therefore could not be used for this analysis.

The human data set was also used for further analysis of the bipolar cell subtypes. The authors already assigned them into 12 well described subtypes (DB1, DB2, DB3a, DB3b, DB4, DB5, DB6, FMB, BB+GB, IMB, RB11, OFF), which we used to subdivide them into ON and OFF, as well as rod bipolar cells.

For a more in depth analysis of amacrine cell subtypes we included an additional data set of Yan et al. [19] (GSE 149715). Again, to start our analysis and processing we used the clusters as published (https://singlecell.broadinstitute.org/single_cell/study/SCP919/). These clusters were evaluated according to their gene expression profile and assigned a neurotransmitter type. As marker genes we used *gad1* and *gad2* (GABA), *slc6a9* and *slc6a5* (glycine), *slc17a8* (glutamate), *chat* (choline) and *Th* (dopamine). If the respective cluster showed a clear expression of the particular marker gene, it was assigned to the corresponding neurotransmitter type. This assignment was not exclusive. A cluster could be assigned to more than one neurotransmitter type. The evaluation was done manually.

Graphs and the preceding analysis were performed using python and in particular the python toolkit scanpy (v1.7.2). The complete code is available from the corresponding author on request.

## Additional information

### Data and code availability statement

The code to analyse the scRNA-seq data sets are available from the corresponding author on reasonable request. data sets used are available online https://singlecell.broadinstitute.org/. Please see the Methods section for further details.

### Author contributions statement

B. P. designed research, performed and analysed computations, and wrote the paper. T. P. performed computations. A. F. designed research, supervised and wrote the paper. All authors have approved the final version of the manuscript and agree to be accountable for all aspects of the work.

